# RNAi screening reveals requirement for PDGFRβ in JEV infection

**DOI:** 10.1101/2021.01.21.427722

**Authors:** Minmin Zhou, Shaobo Wang, Jiao Guo, Yang Liu, Junyuan Cao, Xiaohao Lan, Xiaoying Jia, Bo Zhang, Gengfu Xiao, Wei Wang

## Abstract

Mosquito-borne Japanese encephalitis virus (JEV) causes serious illness worldwide and is associated with high morbidity and mortality. To identify potential host therapeutic targets, a high-throughput receptor tyrosine kinase small interfering RNA library screening was performed with recombinant JEV particles. Platelet-derived growth factor receptor beta (PDGFRβ) was identified as a hit after two rounds of screening. Knockdown of *PDGFRβ* blocked JEV infection, and trans-complementation of PDGFRβ could partly restore its infectivity. The PDGFRβ inhibitor imatinib, which has been approved for the treatment of malignant metastatic cancer, protected mice against JEV-induced lethality by decreasing the viral load in the brain, while abrogating the histopathological changes associated with JEV infection. These findings demonstrated that PDGFRβ is important in viral infection and provided evidence for the potential to develop imatinib as a therapeutic intervention against JEV infection.

## Introduction

Japanese encephalitis virus (JEV) is a major cause of viral encephalitis worldwide, with an estimated 68,000 cases and approximately 13,600 to 20,400 deaths annually (1). JEV belongs to the genus *Flavivirus* in the family *Flaviviridae* and is transmitted between vertebrate hosts by mosquitoes, mainly by *Culex tritaeniorhynchus*. Flaviviruses include other important pathogens such as Zika virus (ZIKV), dengue virus (DENV), West Nile virus (WNV), and yellow fever virus (YFV). Flaviviruses have an approximately 11-kb-long positive-stranded RNA genome containing a single open reading frame (ORF) flanked by untranslated regions (UTRs) at both termini. The ORF encodes three structural proteins, including the capsid (C), membrane (premembrane [prM] and membrane [M]), and envelope (E), and seven nonstructural proteins (2, 3).

E proteins are densely arranged on the surface of the virion and respond to binding and fusion during virus entry into the host cell (4). JEV can infect a plethora of cell types from different species (5). After being released into the skin epidermis by a mosquito bite, JEV spreads from dermal tissues to lymphoid organs, resulting in a transiently mild-to-moderate viremia (6). JEV is neuroinvasive and neurovirulent, and replicates in the central nervous system (CNS) cells such as neurons (7), pericytes (8), astrocytes (9), and microglia (10).

To date, the receptor for JEV is not well characterized. A wide range of cellular surface receptors, as well as attachment factors, have been reported to facilitate JEV entry into different cell types (5). In this study, we focus on the role of receptor tyrosine kinases (RTKs) in JEV infection. RTK has been reported to play critical roles in virus entry and replication (11–15), and RTK inhibitors block multiple steps of the virus life cycle (16–18). Using an RTK RNA interference (RNAi) screen library, we identified that platelet-derived growth factor receptor beta (PDGFRβ) acted as a proviral gene in JEV infection. Moreover, imatinib, a specific PDGFRβ inhibitor, could robustly inhibit JEV infection both *in vitro* and *in vivo*. These results contribute to our understanding of the biology of virus entry and potentially provide new host targets for therapeutic intervention.

## Results

### RTK RNAi library screen identifies *PDGFRβ* as a pro-JEV infection gene

Arrays of four independent small interfering RNAs (siRNAs) targeting 56 RTK genes grouped in a 2 × 2 mix format were transfected into HeLa cells. Forty-eight hours after transfection, the cells were infected with recombinant JEV particles, which were packaged with the structural C, prM, and E proteins and contained a JEV luciferase-reporting replicon (19). Twenty-four hours after infection, the cell lysates were subjected to luciferase assay (Fig. 1A). After the primary screening, eight hits with more than 50% reduction in luciferase activity in both composition pools were selected (Fig. 1B). The synthesized siRNAs with different sequences targeting the eight hits were tested to confirm the primary screening. Only *PDGFRβ* was validated because four synthesized siRNAs in the secondary screen decreased the infectivity to less than 50% (Fig. 1C). The relatively low 12.5% validation rate was due to the exclusion of invalidated siRNAs in secondary analyses and the ruling out of low gene expression with high cellular background qPCR-CT (cycle threshold > 30) values. Among the eight *PDGFRβ* siRNAs, those that diminished the *PDGFRβ* RNA levels inhibited recombinant JEV infection, whereas siRNAs that did not decrease the RNA levels had no effect on infection (Fig. 1C).

**Fig. 1.**
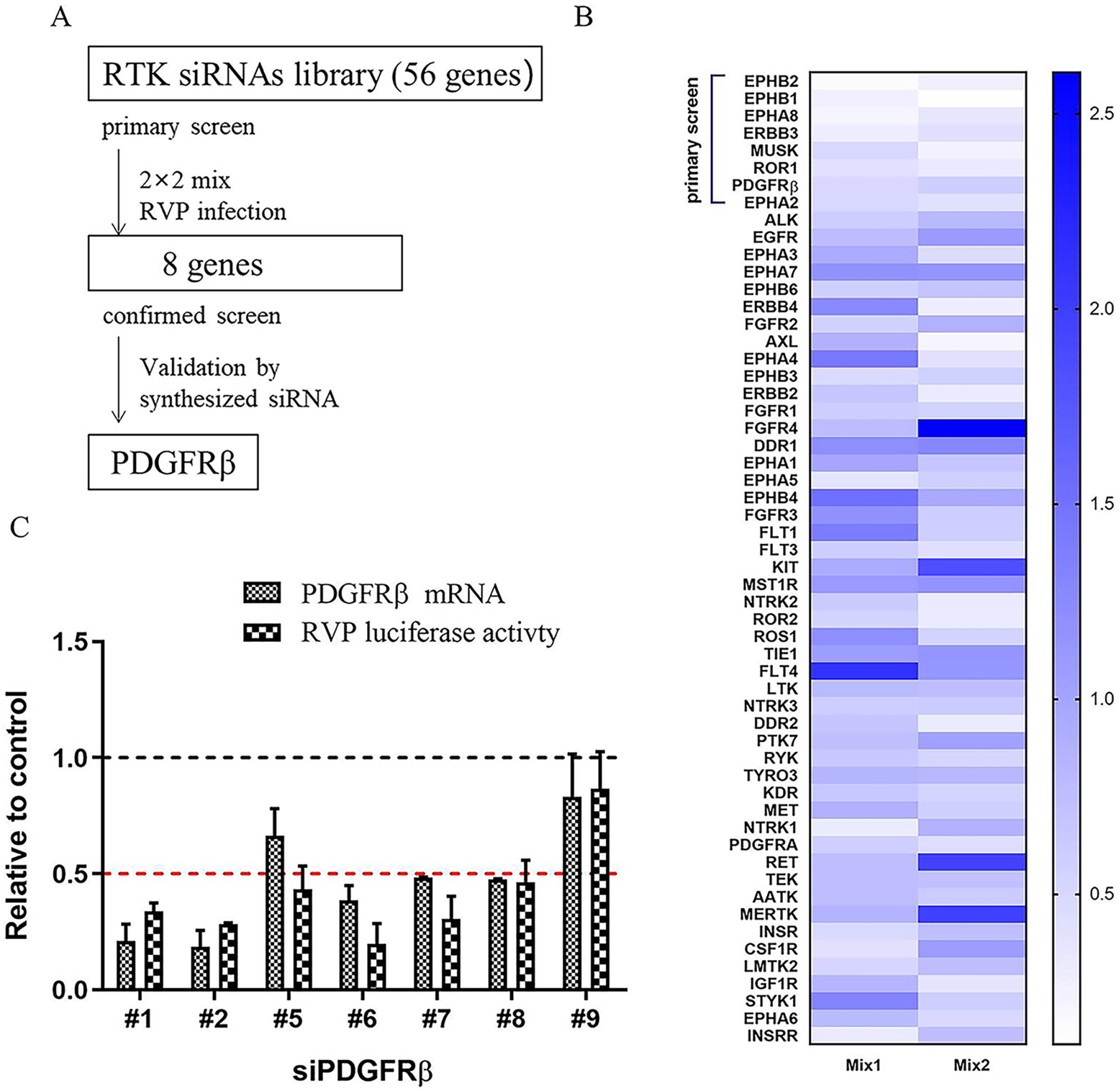
siRNA library screening. (A) Heat map of recombinant JEV infection-induced phenotype. HeLa cells were transfected with an siRNA library targeting 56 RTKs with 2 × 2 mix pools; 48 h later, cells were infected with recombinant JEV particles. Rluc activities were tested 24 h later. The top eight genes were selected because two pools of each gene had an inhibition >50%. (B) siRNA screening assay flowchart. (C) siRNA targeting *PDGFRβ* inhibited recombinant JEV infection. HeLa cells were transfected with siRNAs targeting *PDGFRβ*; 48 h later, cells were infected with recombinant JEV particles. The mRNA level of *PDGFRβ* and Rluc activities were tested 24 h later.

### PDGFRβ is important in JEV infection

To verify the results obtained by the luciferase reporter assays, we investigated the effect of *PDGFRβ* siRNA on authentic virulent JEV AT31 and SA14 strains and the attenuated SA14-14-2 strain. The inhibitory effects were investigated at RNA, protein, and virus production levels using qPCR, Western Blot (WB), and plaque assays, respectively. As expected, sharp decreases in RNA levels of both the virulent and attenuated JEV strains were detected (Fig. 2A, 2D, and 2G). Similarly, expression of the viral non-structural NS3 protein was inhibited by the knockdown of *PDGFRβ* in all the tested JEV strains (Fig. 2B, 2E, and 2H). Intriguingly, the reduction in viral titers was approximately 2 to 3 log units in the JEV AT31 strain (Fig. 2C), and an approximately 1-log-unit decrease was found in the SA14 strain (Fig. 2F), whereas no obvious reduction was detected in the attenuated SA14-14-2 strain (Fig. 2I). Overall, the results presented in Fig. 2 confirmed that PDGFRβ played a critical role in JEV infection and that knockdown of *PDGFRβ* inhibited JEV infection *in vitro*.

**Fig. 2.**
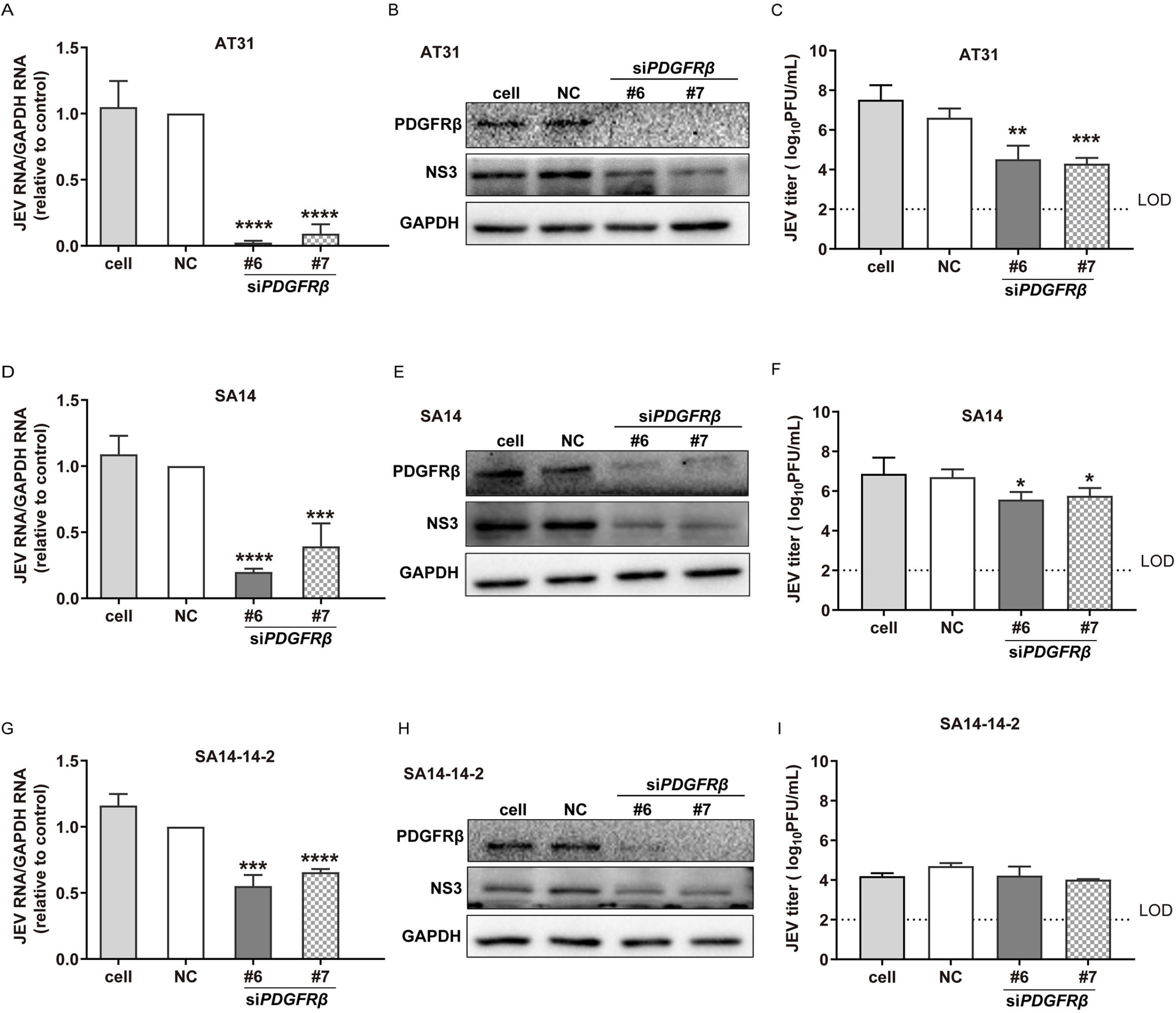
Knockdown of *PDGFRβ* inhibited JEV infection. (A to C) Inhibition against JEV AT31. HeLa cells were transfected with si*PDGFRβ* and negative control, respectively; 24 h later, cells were infected with JEV AT31 (MOI: 0.1). Twenty-four hours later, cell lysates were subjected to qPCR (A) and WB (B), and the supernatants were subjected to plaque assay (C). (D to F) Inhibition against SA14. (G to I) Inhibition against SA14-14-2. Data are presented as means ± SDs from four independent experiments. NC: negative control; LOD: limit of detection. * P < 0.05; *** P < 0.001, **** P < 0.0001.

The effect of PDGFRβ was further confirmed using CRISPR/Cas9 gene editing, by generating a ΔPDGFRβ single-cell clone in HeLa cells and by confirming gene deletion and cell viability (Fig. 3A and 3 B). Infection with JEV AT31 was reduced in ΔPDGFRβ cells (Fig. 3C and 3D), and trans-complementation of PDGFRβ in ΔPDGFRβ HeLa cells restored infectivity in a dose-dependent manner (Fig. 3E).

**Fig. 3.**
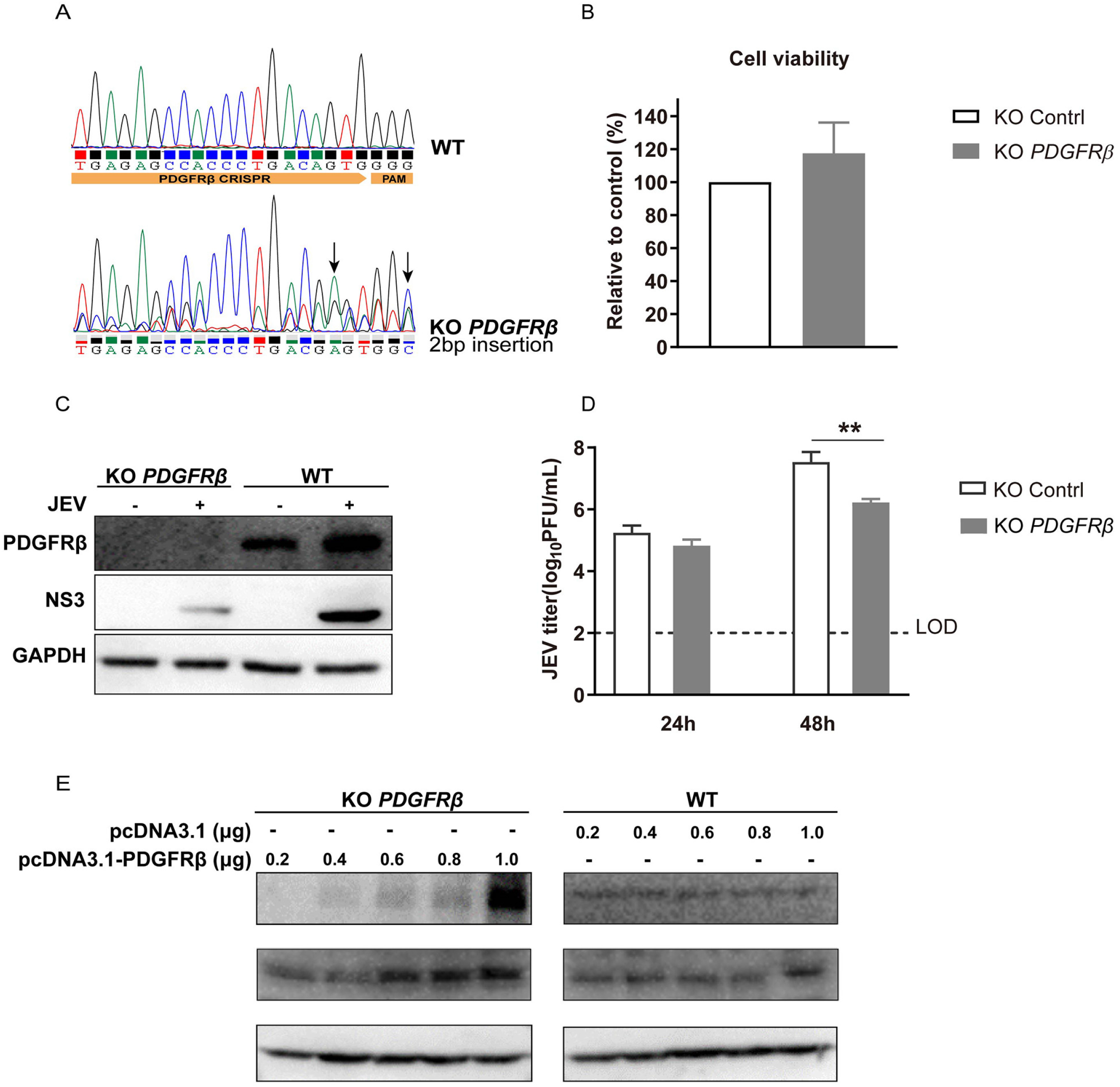
Validation of the inhibitory activity using knockout (KO) cell line. (A) Sequencing chromatogram of the WT and PDGFRβ KO cells. (B) PDGFRβ KO showed little effect on cell viability. The viabilities of WT and PDGFRβ KO HeLa cells were assessed using a luminescent cell viability assay kit. (C) WB assay. WT and PDGFRβ KO cells were infected with JEV AT31 (MOI: 0.1). Cell lysates were subjected to WB assay 24 h later. (D) Plaque assay. WT and PDGFRβ KO cells were infected with JEV AT31 (MOI: 0.1). The supernatants were subjected to plaque assay 24 h and 48 h later. Data are presented as means ± SDs from three independent experiments. LOD: limit of detection. ** P < 0.01. (E) Trans-complementation of *PDGFRβ* in ΔPDGFRβ HeLa cells restored infectivity. ΔPDGFRβ or control HeLa cells were inoculated with JEV and subjected to WB for detection of PDGFRβ, JEV NS3, and GAPDH.

### Imatinib blocks JEV infection *in vitro*

Imatinib is a tyrosine kinase inhibitor that specifically targets the kinase activity of PDGFRβ, ABL, and c-KIT and has been approved for the treatment of chronic myeloid leukemia and gastrointestinal stromal tumors (20–24). As PDGFRβ is important in JEV infection, we further tested whether imatinib could inhibit JEV infection. Four cell types were used to evaluate the antiviral effect. As shown in Fig. 4A, imatinib inhibited JEV AT31 infection in Baby hamster kidney (BHK-21) cells, African green monkey kidney (Vero) cells, human cervix epithelial (HeLa) cells, and human liver epithelial-like (Huh7) cells, with IC_50_ values ranging from 4.987 to 26.17 μM (Fig. 4A). Notably, 50 μM imatinib showed mild to little effect on the viability of all the tested cell types (Fig. 4B).

**Fig. 4.**
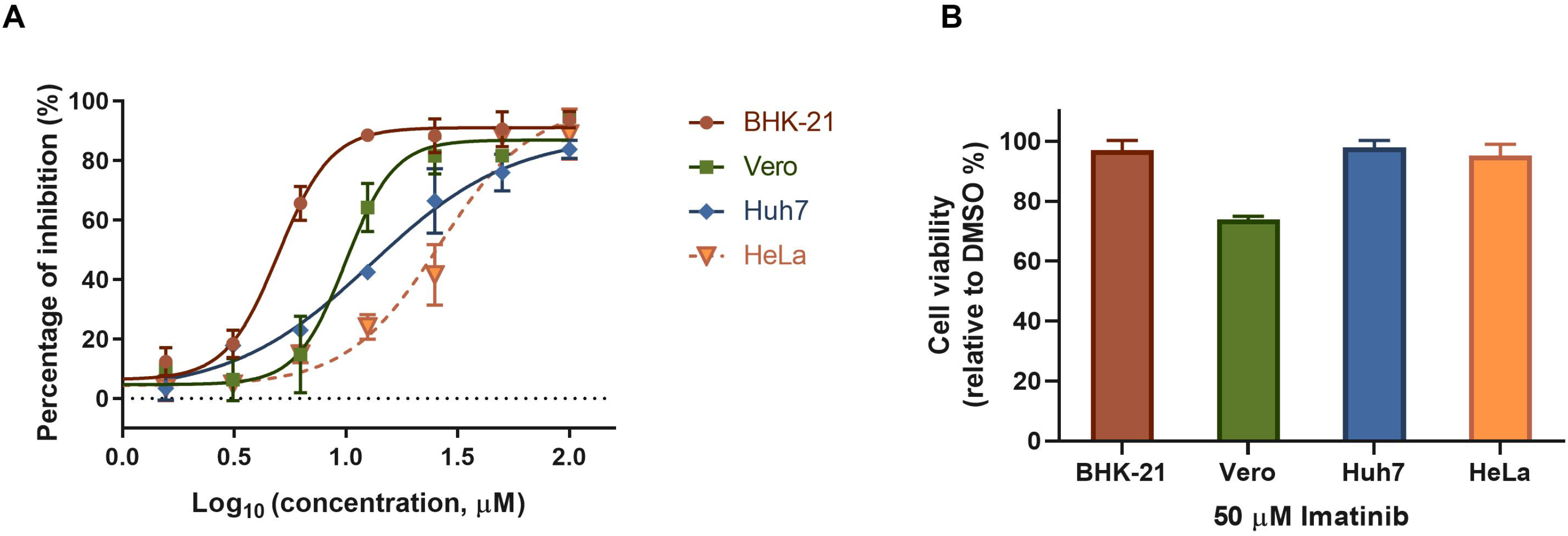
Imatinib blocked JEV infection in different cell types. (A) Dose-response curves of imatinib for inhibition of JEV infection on different cell types. (B) Cells were incubated with 50 μM imatinib for 24 h, and the cell viabilities were evaluated using CCK8 assay.

### Imatinib inhibits JEV infection *in vivo*

As imatinib exhibited a robust inhibitory effect on JEV infection, we further examined the protective effect of imatinib against JEV-induced lethality in a mouse model. As expected, JEV-infected mice began to show symptoms, including limb paralysis, restriction of movement, piloerection, body stiffening, and whole-body tremor, from day 3 post-infection. Within 21 days post-infection, eight mice in the JEV-infected group (n =10) succumbed to the infection, with the mortality rate being 80%. Whereas imatinib treatment delayed the disease onset and reduced the mortality rate to 10% (n =10) (Fig. 5A). Moreover, mice in the imatinib-treated group showed slightly abnormal behavior, similar to the findings for the mice in the imatinib alone group. These results indicated that imatinib provided effective protection against JEV-induced mortality.

**Fig. 5.**
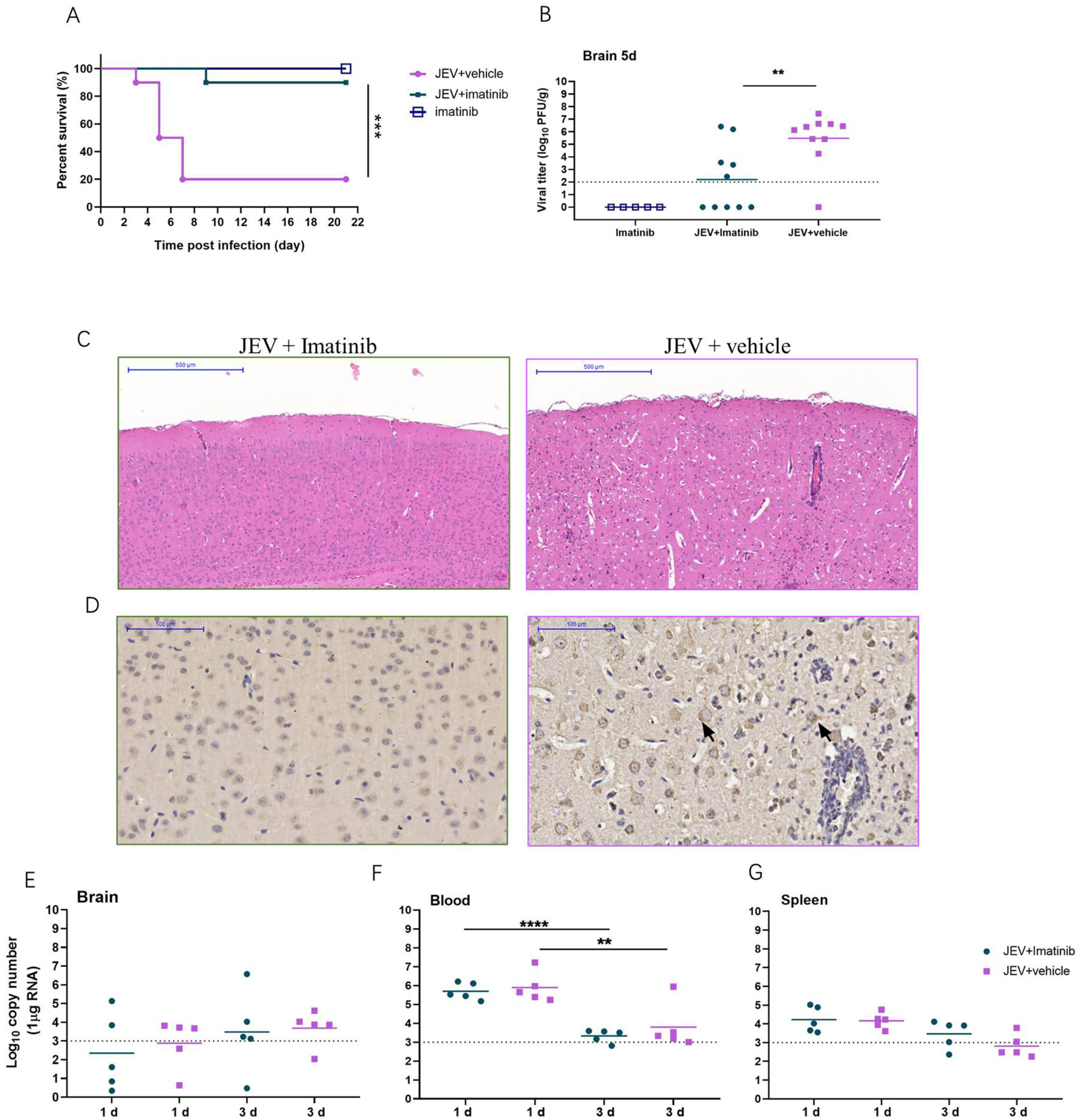
Imatinib protected mice from JEV infection. C3H/He mice were infected with 1 × 10^7^ PFU of JEV together with imatinib (30 mg/kg, i.p. once a day). (A) Survival curve for either group was monitored for five days (n = 10). (B) The viral loads in mouse brains were measured by plaque assay on day 5. (C) Histopathological analysis of the mouse brain from imatinib and vehicle treatment group. Arrows indicate the histopathological changes such as meningitis, perivascular cuffing, and glial nodules. Bar: 500 μM. (D) IHC analysis of expression of prM protein in mouse brain. Staining for JEV prM protein. Imatinib treatment alleviated the histopathological changes in mice caused by JEV infection. Imatinib treatment decreased the viral loads in mice brain. Bar: 100 μM. (E-G) The viral loads in brain (E), blood (F), and spleen (G) were measured by qRT-PCR on days 1 and 3, respectively. Dashed lines indicate limit of detection. **** P < 0.0001; ** P < 0.01.

To further relate these protective effects to the viral load and histopathological changes in the mouse brains, the viral titer was determined, and mouse brain sections were collected and assayed at day 5 post-infection. As shown in Fig. 5B, imatinib treatment significantly reduced the viral load in infected mice compared to no treatment (P = 0.0032), and no plaques formed in the imatinib group (Fig. 5B). Similarly, apparent damage to the brain, including meningitis, perivascular cuffing, vacuolar degeneration, and glial nodules, was observed in the JEV-infected group on day 5 post-infection, while imatinib treatment remarkably alleviated these phenomena (Fig. 5C). Immunohistochemical (IHC) analysis revealed diffuse and coalescing prM-staining density in neurons in the JEV-infected group, whereas it was absent in the imatinib-treated group. These results indicated that the alleviation of histopathological changes was accompanied by a reduction in the viral load as well as a reduction in the rate of mortality, further confirming the potential curative effects of imatinib on viral encephalitis.

As JEV was rapidly cleared from the blood after inoculation and was present in the lymphatic system during the preclinical phase (25–28), the effects of imatinib on infection of the brain, blood, and spleen were evaluated at earlier time points (day 1 and 3 post-infection) to detect whether the drug reduced the peripheral viral loads. As shown in Fig. 5D to 5E, imatinib had little effect on peripheral JEV infection, which confirmed that imatinib protected the mice against JEV-induced lethality by reducing the viral load in the brain.

## Discussion

RTKs are high-affinity cell surface receptors for many polypeptide growth factors, cytokines, and hormones (29, 30). Generally, ligand binding activates RTKs by inducing receptor dimerization and autophosphorylation of their own kinase domains, thus leading to the activation of intracellular signaling networks (31). Several RTKs have been reported to play essential roles in viral infection. Among these, the TAM subfamily (Tyro3, Axl, and Mertk) has been postulated to be an entry receptor for flaviviruses (32, 33). However, it was recently reported that TAM is dispensable for ZIKV infection, as ZIKV was able to infect and replicate in TAM receptor knockout (KO) mice (34). Intriguingly, WNV and La Crosse virus infections of neurons can be promoted in mice lacking Axl and Mertk (35). Similarly, the neuroinvasion of JEV was enhanced in mice lacking Axl (36). In this study, we attempted to identify the critical RTK molecules involved in JEV infection. In the first siRNA library screening, knockdown of either *Tyro3* or *Axl* led to mild reduction of recombinant JEV infection with inhibition of <50%, whereas knockdown of *Mertk* slightly promoted recombinant JEV infection. Among the eight primary hits, only *PDGFRβ* was validated with additional siRNAs. Notably, screening was carried out on cell-based high throughout screening and validation, which might be different from natural infection *in vivo*. As described above, the absence of Axl and Mertk would increase blood-brain barrier permeability, thus enhancing the infectivity of WNV (35). Similarly, deficiency of Axl enhances the production of IL-1α and thus accelerates the neuroinvasion of JEV (36). It is impossible to establish *PDGFRβ* KO mice because deletion of this gene is lethal (37). Whether PDGFRβ serves as a potential receptor of JEV, or PDGFRβ recruits the intracellular signal network to act as a proviral/antiviral protein, or whether the contribution of PDGFRβ to inflammation affects the infectivity of JEV needs to be further investigated.

However, the validation of PDGFRβ still provides a promising target for the development of anti-JEV therapy. To this end, we tested the efficacy of imatinib against JEV infection both *in vitro* and *in vivo*. The pharmacokinetic and pharmacodynamic parameters of imatinib have been well characterized (38). The maximum plasma concentrations (Cmax) of imatinib at steady state in patients with gastrointestinal stromal tumors (GISTs) and chronic myelogenous leukemia (CML) have been identified as 2.9 μg/mL (4.1 μM) and 2.3 μg/mL (3.2 μM), respectively (22, 39), which was comparable to the IC_50_ reported in this study. It has been reported that intraperitoneal (i.p.) administration of 50 mg/kg/day imatinib to mice for 22 consecutive days would lead to local toxicity at the i.p. injection site, whereas little organ toxicity was observed (40). Based on our experience, when re-purposing a drug used to treat chronic diseases to combat infectious diseases, it is generally necessary to raise the drug dose to an extremely high level (27, 28, 41, 42). Intriguingly, the dose used in this *in vivo* study (30 mg/kg/day, i.p.) was relatively close to the dose used in the clinic for the treatment of GIST (400–600 mg/day, p.o.) and CML (400– 800 mg/day, p.o.) (22, 38), suggesting that imatinib may be a potential safe treatment for JEV infection. Our finding that PDGFRβ is important for JEV infection and the approved drug imatinib blocks the infection suggest that an effective re-purposing of PDGFRβ inhibitor is a possible treatment strategy for JEV infection.

## Materials and Methods

### Cells and virus

BHK-21 cells, Vero cells, HeLa cells, and Huh-7 cells were cultured in Dulbecco’s modified Eagle’s medium (DMEM; HyClone, Logan, UT, USA) supplemented with 10% fetal bovine serum (GIBCO, Grand Island, NY, USA).

JEV AT31 and SA14 strains were generated using the infectious clones of pMWJEAT-AT31 (kindly provided by T. Wakita, Tokyo Metropolitan Institute for Neuroscience) and pACYC-JEV-SA14 (GenBank accession no. U14163), respectively, as previously described (19). Briefly, the infectious clone plasmids were linearized, subjected to *in vitro* transcription, and delivered to BHK-21 cells via electroporation. Three days later, the supernatant was collected, propagated, and titrated in BHK-21 cells. The SA14-14-2 strain is a commercial vaccine obtained from Keqian Biology (Wuhan, China). JEV recombinant replicon particles were generated as previously described (4). Briefly, the JEV luciferase-reporting replicon (SA14/U14163-Replicon) was linearized, subjected to *in vitro* transcription, and delivered into BHK-21 cells via electroporation. After 72 h, the cells were transfected with pCAGGS-CprME, and the supernatant was collected 48 h later.

### RTK siRNA library screening

The RTK RNAi library contained siRNAs targeting 56 human RTK genes (Qiagen, Dusseldorf, Germany), which included four different pairs of siRNAs for each gene, adding up to 226 pairs of siRNAs in total. Two × two pooled siRNAs were transfected into pre-seeded cells. Forty-eight hours later, the cells were infected with recombinant JEV particles. The cell lysates were subjected to luciferase activity testing (Promega, Madison, WI, USA) 24 h later. Primary hits were identified as those exhibiting a more than 50% decrease in luciferase activity relative to the control in the two different composition pools. Additional *PDGFRβ* siRNAs (Table 1, synthesized by Gene Pharma, Suzhou, China) were used for knockdown assays with recombinant JEV particles.

**Table 1.**
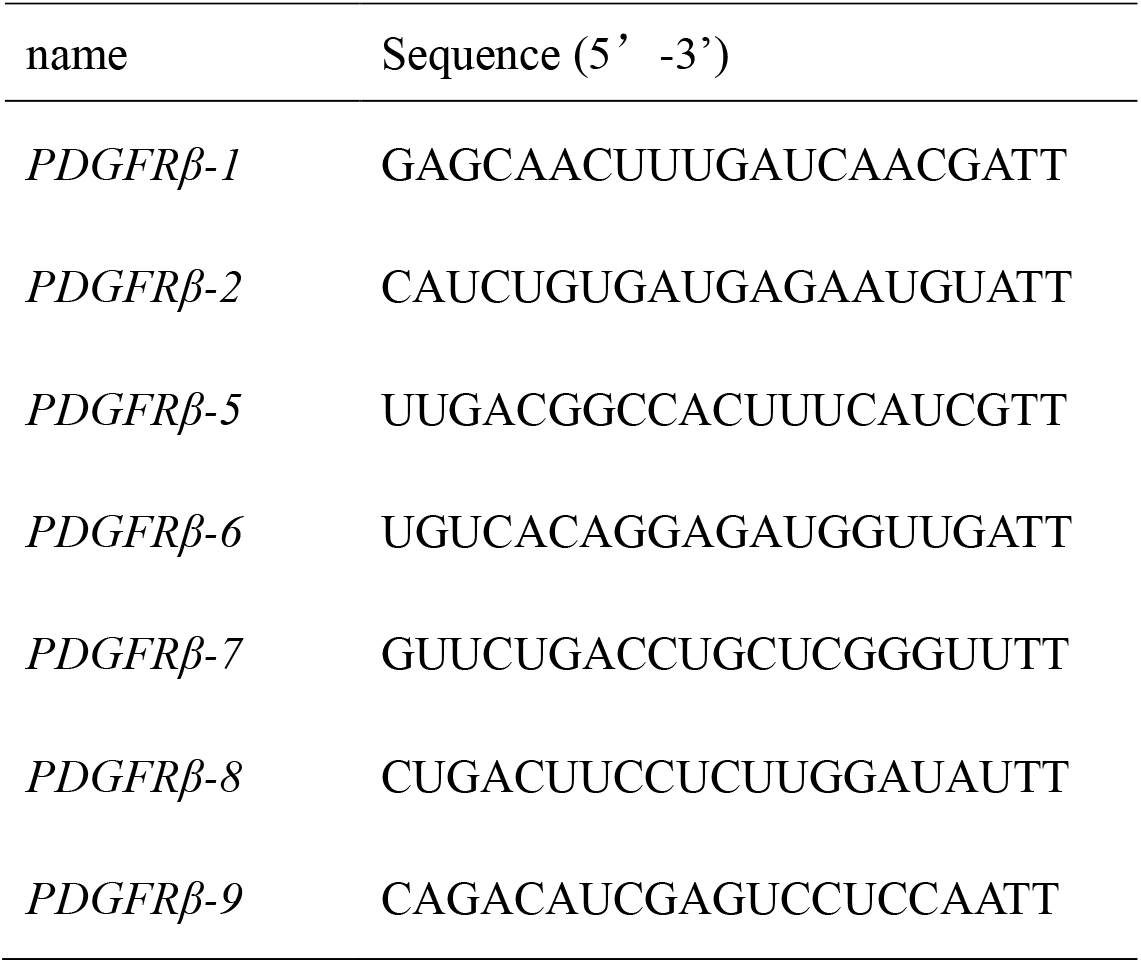
Sequences of siRNAs used in confirmed screen.

### Cell viability

CellTiter-Lumi™ Luminescent Cell Viability Assay Kit (Beyotime, Shanghai, China) was used to evaluate the cell viability of the KO cell line. Cell counting kit-8 (CCK-8) (GlpBio Technology, Montclair, CA, USA) assay was carried out to evaluate the effect of imatinib on cell viability. Briefly, imatinib at the indicated concentrations was added to pre-seeded BHK-21, Vero, HeLa, and Huh-7 cells in 96-well plates. Twenty-four hours later, cell viability was measured using the CCK-8 kit, and the light absorption value was read at 450 nm.

### Antiviral assay

Cells in 96-well plates were infected with JEV AT31 (MOI: 0.1) for 1 h. Imatinib at the indicated concentrations was added to the cells from 1 h pre-infection to 23 h post-infection. The antiviral effects of imatinib were evaluated using an Operetta high-content imaging system (PerkinElmer) (28). Briefly, cells were fixed with 4% paraformaldehyde. Cells were then permeabilized using PBS with 0.2% Triton X-100 for 15 min and blocked with 5% FBS (Gibco), followed by treatment with the primary anti-JEV prM antibody (4) and staining with DyLight 488-labeled rabbit IgG antibody (KPL, Gaithersburg, MD, USA). Nuclei were stained with 4’,6-diamidino-2-phenylindole (DAPI, Sigma-Aldrich, St. Louis, MO, USA). Nine fields per well were imaged, and the percentages of infected and DAPI-positive cells were calculated using the associated the Harmony 3.5 software.

The effect of PDGFRβ on JEV infection was evaluated using qPCR (43), WB assay, and plaque assay. The antibodies used in the WB assay included anti-PDGFRβ rabbit mAb (1:1000, Cell Signaling Technology, Danvers, CA, USA), anti-JEV NS3 rabbit antiserum (kindly provided by C.J. Chen, Taichung Veterans General hospital, 1:1000), anti-GAPDH mouse mAb (ABclonal, Wuhan, China; 1:1000), horseradish peroxidase (HRP)-linked goat anti-rabbit IgG, and HRP-linked goat anti-mouse IgG (Proteintech, CHI, USA, 1:5000).

### Establishing the ΔPDGFRβ cell line

*PDGFRβ*-targeted sgRNA (5’-CCGGTGAGAGCCACCCTGACAGTG-3’) was cloned into the pGuide-it-ZsGreen1 vector (Takara, Biomedical Technology, Beijing, China). This plasmid could simultaneously express *Cas9*, the *PDGFRβ*-targeting sgRNA, and the bright green fluorescent protein. HeLa cells were transfected with 2 μg plasmid per well in 6-well plates, and the cells were collected 24 h after transfection. Bright green fluorescent expressing monoclonal cells were sorted by flow cytometry and proliferated to form a cell population after nearly two weeks of culture. The KO cell line was validated by sequencing with the primers of f-5’-TTGTTAAAAGGGAAGATTAGCAAGT-3’ and r-5’-AAAGGTAAGGAAAAGGGACCATTTA-3’.

### Imatinib administration to JEV-infected mice

All animal experimental procedures were carried out according to ethical guidelines and were approved by the Animal Care Committee of the Wuhan Institute of Virology (permit number, WIVA25201901).

Adult C3H/He female mice (4 weeks old) (44, 45) were randomly divided into three groups (30 to 36 mice each): JEV-infected, imatinib-treatment, and imatinib alone. Mice in the JEV-infected group were administered i.p. injection with 10^7^ PFU of JEV strain AT31 in 100 μL PBS, those in the imatinib group were treated with 30 mg/kg imatinib once a day for 21 consecutive days, and those in the imatinib-treated group were treated with 10^7^ PFU of JEV strain AT31 on the first day along with 30 mg/kg imatinib once a day for 21 consecutive days. Ten mice in each group were monitored daily to assess behavior and mortality. The remaining mice were sacrificed on days 1, 3, 5, and 21 post-infection, and the brain, liver, and spleen were harvested for analysis.

To detect the viral burden using plaque assay, brain tissues were dissected, ground with 300 μL PBS, and the supernatants containing virus particles after centrifugation were serially diluted 10-fold prior to infecting BHK-21 cells. Viral titers were assessed per gram of tissue using a plaque assay.

To detect the viral burden by qPCR, tissue samples and blood were extracted with the RNAprep pure tissue kit (catalog no. DP431, Tiangen, Beijing, China) and RNAprep pure blood kit (Tiangen; catalog no. DP433), respectively. The viral burden was calculated on a standard curve produced using serial 10-fold dilutions of plasmids carrying the infectious clone.

For histopathological analysis, brain samples were sectioned and stained with hematoxylin and eosin (Goodbio Technology Co., Wuhan, China). For IHC staining, sections were incubated sequentially with primary anti-JEV prM antibodies (dilution, 1:200) and HRP-conjugated secondary antibodies. Additionally, 3,3’-diaminobenzidine (DAB) was used for color development. Pathological changes and IHC staining were analyzed using the Pannoramic Viewer Plus software (3DHISTECH, Budapest, Hungary).

### Statistical analysis

Statistical analysis was performed using GraphPad Prism 8 software (GraphPad Inc., La Jolla, CA, USA). Survival data were analyzed using the log-rank test and all other statistical analyses were assessed with the Student’s t-test.

## Acknowledgements

We thank the Center for Instrumental Analysis and Metrology, and Core Facility and Technical Support, Wuhan Institute of Virology, for providing technical assistance. This work was supported by the National Key Research and Development Program of China (2018YFA0507204), the National Natural Sciences Foundation of China (31670165), Wuhan National Biosafety Laboratory, Chinese Academy of Sciences Advanced Customer Cultivation Project (2019ACCP-MS03), the Open Research Fund Program of the State Key Laboratory of Virology of China (2018IOV001).

